# *In situ* cryo-electron tomography reveals local cellular machineries for axon branch development

**DOI:** 10.1101/2021.07.28.454169

**Authors:** Hana Nedozralova, Nirakar Basnet, Iosune Ibiricu, Satish Bodakuntla, Christian Biertümpfel, Naoko Mizuno

## Abstract

Neurons are highly polarized cells forming an intricate network of dendrites and axons. They are shaped by the dynamic reorganization of cytoskeleton components and cellular organelles. Axon branching allows to form new paths and increases circuit complexity. However, our understanding of branch formation is sparse due to technical limitations. Using *in situ* cellular cryo-electron tomography on primary mouse neurons, we directly visualized the remodeling of organelles and cytoskeleton structures at axon branches. Strikingly, branched areas functioned as hotspots concentrating organelles to support dynamic activities. Unaligned actin filaments assembled at the base of premature branches and remained while filopodia diminished. Microtubules and ER co-migrated into preformed branches to support outgrowth together with accumulating compact ~500 nm mitochondria and locally clustered ribosomes. We obtained a roadmap of events and present the first direct evidence of local protein synthesis selectively taking place at axon branches, allowing to serve as unique control hubs for axon development and downstream neural network formation.

## Introduction

The development of neurons with their extremely polarized structure and function is unique. Several long protrusions or neurites extend from the main cell body, the soma, where the nucleus is located. One of the protrusions develops into the axon, while the remaining protrusions develop into dendrites(Barnes and Polleux, 2009). Axons are functionally distinct from dendrites; dendrites receive signals from the axons of upstream cells via synaptic connections, whereas axons transmit signals to the dendrites of downstream cells. The molecular organization of the axon is uniquely suited to support local developmental processes, reflecting its specialized function. Within the backbone of the axon, microtubules form parallel bundles with their plus ends oriented towards the distal end of the axon(Baas et al., 1988; Baas and Lin, 2011; Stepanova et al., 2003; van Beuningen and Hoogenraad, 2016). An actin-rich growth cone at the tip of a growing axon probes extracellular signalling molecules to identify the synaptic target on dendrites of an adjacent neuron (Dent and Gertler, 2003; Lowery and Van Vactor, 2009).

The long distance from the soma to the tip of an axon indicates that there is a regulatory system controlling local molecules(Dalla Costa et al., 2021; Goldberg, 2003; Holt et al., 2019; Stiess et al., 2010). This regulatory system includes differential expression of and enrichment for proteins that are critical for the local development and regulation of axonal homeostasis. In particular, clusters of mRNAs localized within dendrites and axons(Cioni et al., 2018; Holt et al., 2019; Poon et al., 2006; Taylor et al., 2009) indicate local protein synthesis. The types of locally enriched mRNAs depend on the developmental stage and their location within a neuron(Cioni et al., 2018). During the growth phase, mRNAs coding for proteins of the synthesis machinery such as ribosome and for cytoskeletal components, which are needed to extend the cell, are found predominantly in the growth cones(Bassell et al., 1998; Zivraj et al., 2010). This type of regulation has also been observed in distal axons during regeneration(Gumy et al., 2010). However, despite the critical role of local translation in axons, actions of the translation process is not well understood. While there has been growing evidence for the presence of ribosomes along the axons (Koenig et al., 2000; Noma et al., 2017; Tcherkezian et al., 2010), there is no direct observation for protein synthesis that is accompanied by cytoskeleton and organelle re-organization and this has been a major challenge. Direct observations of the local axonal environment at a molecular level would aid in our understanding of neuronal growth and local reorganization.

During the development of the nervous system, axon branching serves to propagate signals to diverse regions of the nervous system(Kalil and Dent, 2014). Axon branching begins with the formation of actin-rich filopodia, short cellular protrusions, resulting from a signaling pathway that is induced by extracellular cues (Spillane et al., 2013; Tang and Kalil, 2005; Wang et al., 1999). Filopodia are the structural precursors of axon branches, and they develop to mature branches by the action of microtubules recruitment to the filopodia(Dent and Kalil, 2001; Gallo, 2011; Gallo, 2013). At the axon branch point, there is an enrichment of mRNAs(Spillane et al., 2013) encoding proteins such as beta-actin(Wong et al., 2017) that may be required to form the initial premature branched axon(Donnelly et al., 2013). Interestingly, it has also been reported that mitochondria are enriched at axon branching points(Spillane et al., 2013), which may provide energy (Sheng, 2017) and may adjust the Ca^2+^ concentration for signal transduction and branch morphogenesis(Hutchins and Kalil, 2008). While the information for those individual components are available, the orchestration that control the organized assembly of the protein synthesis machinery, organelles, and the cytoskeleton at axon branches are largely unknown.

To understand the organization of the key players for axon branching, we directly visualized the molecular organization of both premature and mature axon branching sites of mouse primary neurons by cryo-electron tomography (cryo-ET). We show the localization of small, ~500 nm mitochondria and short actin fragments at the branches. An intricate network of endoplasmic reticulum (ER) membranes was often found between microtubule bundles and mitochondria, with occasional interactions of the ER membrane with microtubules and mitochondria. The ER was generally accompanied by microtubules at the mature axon branch, indicating that ER migration is guided exclusively by microtubules. We further demonstrate the first direct observation of clusters of ribosomes accumulated preferentially at axon branches. In some cases, the ribosome clusters attached to meshed-planar ER membranes as ER tubes widened, spreading into the space made for the branching activity. Subtomogram-averaging and distance analysis indicated that the clustered ribosomes formed active polysomes. Isolated ribosomes, possibly monosomes (Su et al., 2016) synthesizing a small number of proteins, were also seen but only sparsely. This is in stark contrast to ribosomes found at synapses, where the majority of ribosome formed monosomes (Biever et al., 2020), highlighting the requirement for different types and amounts of newly synthesized proteins at various local environments within an axon. Our observations provide a comprehensive picture of the axon branching process.

## Results

### Structural analysis of branching axons

To visualize the molecular organization of axons and axon branches, we prepared primary neuronal cell cultures from hippocampus and thalamus explants of mouse embryos at stage E15.5. We observed 117 tomographic reconstructions of axons (Table 1) corresponding to ~260 μm in total length. Among these reconstructions, 43 axon branches contained microtubules in the branch site, indicating mature axon branches (a representative tomogram in Fig. 1, Fig. 2G, 2H, movie 1), and 20 nascent branch points had membrane protrusions made of actin-containing filopodia but lacked microtubule-reinforcements, indicating premature axon branches (representative tomogram in Fig. 2, movie 2). At mature branches, microtubules were tightly packed along the axon (Fig. 1B and C, arrows, also Fig. S6C, S6D [axon shaft]) but were looser at the branch points where sections of microtubules spread apart to enter the branch. In mature branches, short unorganized fragments of actin appeared to fill the space (Fig. 1C, light blue), which lacked other large organelles. In contrast, in premature branches, a dense parallel actin network formed filipodia perpendicular to the axis of the axon, followed by accumulation of short fragments of actins at the connection to the axon, while microtubules ran along the axon (Fig. 2C).

**Table 1.** Numeric summary of analysed tomograms. Showing localization of tomographic data collection site within axon and individual cellular features observed at the given locality.

| *location* | *total number* | *with mitochondria* | *with ribosome clusters* | *with ribosomes + mitochondria* | *with ER present inside branch* |
| --- | --- | --- | --- | --- | --- |
| mature branch | 43 | 18 | 16 | 9 | 41 |
| premature branch | 20 | 5 | 13 | 5 | 3 |
| branch total | 63 | 23 | 29 | 14 | 44 |
| other | 54 | 9 | 13 * | 2 | - |
| total | 117 | 32 | 42 | 16 | - |

\* at filopodium / growth cone = 11, varicosity = 2, tight shaft = 0

**Figure 1.**
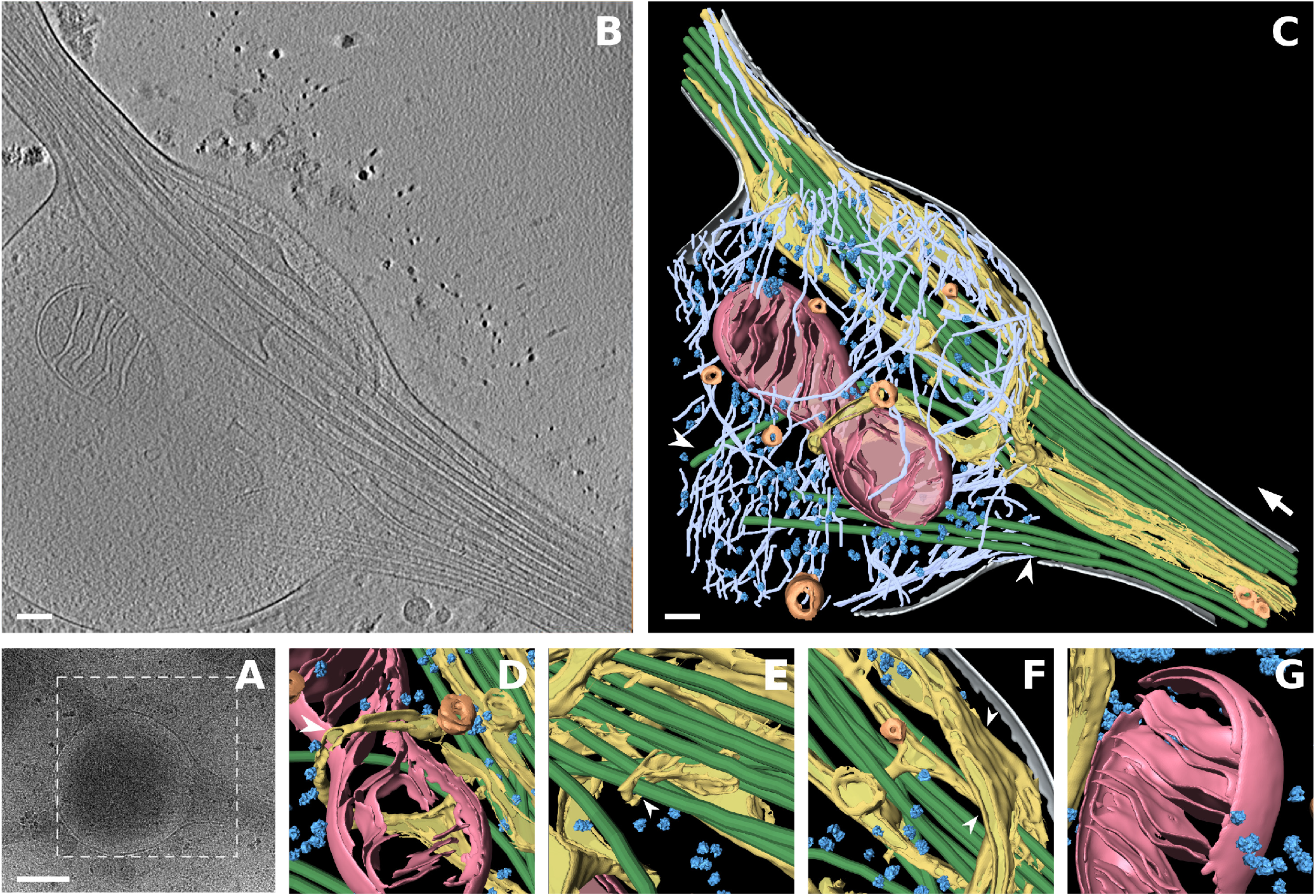
Cryo-electron tomogram of mature axon branch. A) Low magnification view at a mature axon branching site. White box depicts area of tomographic data collection shown in B. B) A slice of the tomogram reconstruction of the branching site. C) Segmentation of the 430-nm thickness tomographic volume from B. Colour code: grey – cellular membrane, green – microtubules, light blue – actin, pink – mitochondria, yellow – ER, dark blue – ribosomes, orange – vesicles, White arrow shows direction of bundled microtubules following axon growth, white arrowheads depict microtubules entering the branch. D)-G) Zoom in views of segmented volume, D) ER wrapping around mitochondrion (white arrowhead), E) ER wraps around microtubule, F) ER forms a flat sheet (white arrowhead), G) ribosomes in the vicinity of mitochondrion. Scale bars: A, 500 nm; B, C 100 nm.

**Figure 2.**
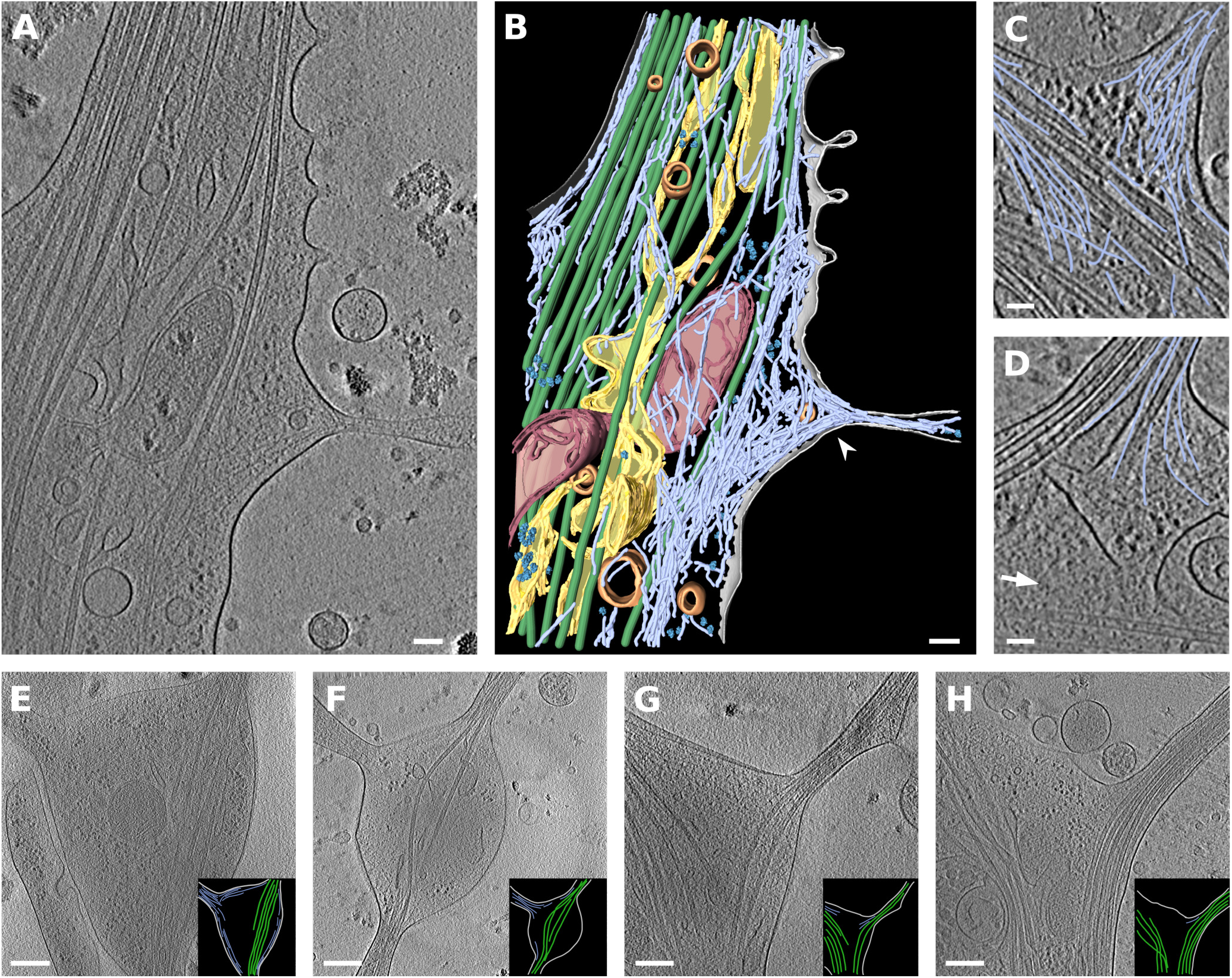
Cryo-electron tomogram of premature axon branch. A) Slice from tomogram reconstruction of premature branch. B) 353-nm segmented volume of tomogram A. Color code – cellular membrane – grey, microtubules – green, mitochondria - pink, ER – yellow, ribosomes – dark blue, vesicles – orange, actin – light blue. White arrowhead depicts filopodium filled with actin. C) and D) An example of actin arrangement in branch, tomographic slice with traced actin (light blue). C) Actin in the filopodia of premature branch. D) Actin in mature branch, white arrow shows direction of main axon growth. E)-H) A slices from axon branch tomograms. E) and F) premature branches, G) and H) mature branches. Black inserts depict the traced cell membrane – grey, microtubules – green and actin – light blue. Scale bars: A,B 100 nm, C,D 50 nm, E-H – 250 nm.

### Organelle organization within axon branches

In addition to the unique organization of the cytoskeleton, branching sites showed localization of mitochondria. Out of the surveyed areas including 63 branches (43 mature and 20 premature) and 54 unbranched areas (44 axon shafts, 10 growth cones), we found 44 mitochondria among 23 mature and premature branches, with only 9 mitochondria along axon shafts, indicating a strong association of mitochondria with axon branches. Axon branch mitochondria had a median size of 500 nm (N=44) (Fig. 3A, 3B) This is in accordance with previous studies showing that axonal mitochondria are significantly smaller than dendritic mitochondria, which are about 4–13 μm (Delgado et al., 2019; Lewis et al., 2018; Popov et al., 2005). Mitochondrial size may be related to the local Ca^2+^ concentration or to presynaptic neurotransmitter release (Lewis et al., 2018), which then eventually lead to terminal axon branch formation. Branching axons accumulated several adjacent ~500 nm mitochondria and mitochondria undergoing fission instead of a single large mitochondrion (Fig. 3A, 1D, S1,).

**Figure 3.**
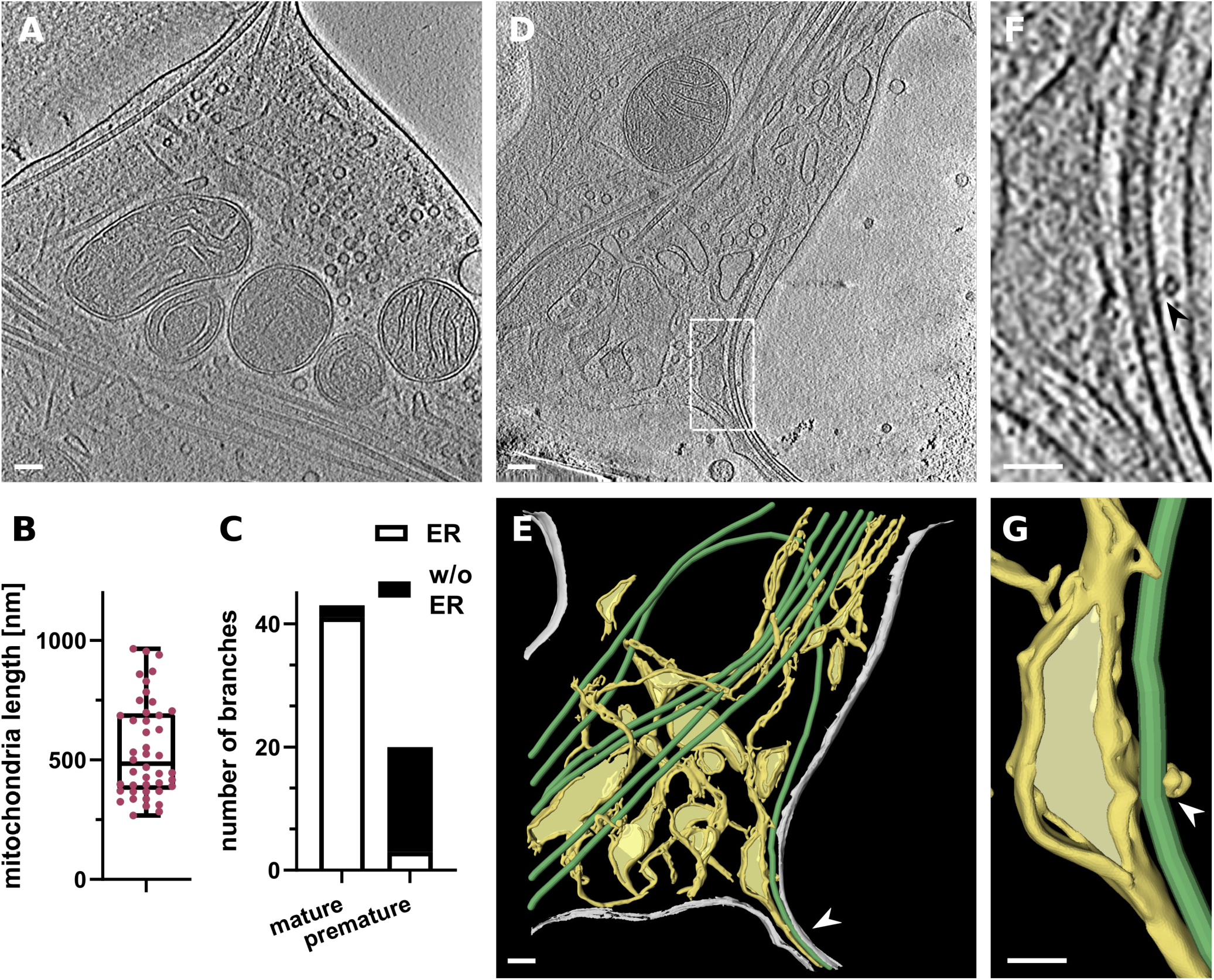
Mitochondria and ER in axon branch. A) Tomographic slice of mitochondria enriched in axon branching site. B) Analysis of mitochondria length. Median length: 500.1 nm N=44. C) Bar graph depicting number of branches with and without ER in daughter branch for mature branches with MT and for premature branches with only actin. D) Slice from tomogram showing different kind of ER – thin tubes and flat sheets. White square depicts area in F and G. E) Segmented tomogram D, color code: grey – cell membrane, green – microtubules, yellow –ER, white arrowhead shows ER in branch together with microtubules. F) ER – MT contact (black arrowhead). G) Segmentation of ER tube wrapping around microtubule (white arrowhead). Scale bar: A,D,E 100 nm, F,G = 50 nm.

The small mitochondria were often next to the ER network, and tubular ER was occasionally wrapped around the mitochondria, either loosely (Fig. 1D), or more tightly contacting to the wide surfaces of their membranes (Fig. 2A, 2B, S1A). The wrapping of the mitochondrion by the ER likely represents a stage of ER-facilitated fission of mitochondria(Lewis et al., 2016; Wu et al., 2017), but it appears the fission of mitochondria may not require wrapping by the ER (Fig. S1B, S1D). The ER at branching sites took on a planar mesh-like spreading form (Fig. 3B, 3D), while along the axon shaft it adopted a thin, tightly-packed tubular structure. The size of the ER ranged from thin tubes as small as 4 nm (Fig. S2A, S2D) to wide flat areas over 200 nm in width (Fig. 1F, 3D-G, 5G). ER membranes were often intertwined with microtubules (Fig. 1E and F, S2B) or occasionally tethered to the walls of microtubules (Fig. 3F, 3G, S2C). We found no density bridging microtubules and ER membranes, presumably due to the low signal-to-noise ratio, though ER-microtubule connections can be made by molecules like p180, CLIMP63, and kinectin(Cui-Wang et al., 2012; Farias et al., 2019; Shibata et al., 2010). Interestingly, we found that the ER propagation to axon branches (Fig. S3) occurred for 41 mature branches out of 43, but for only 3 out of 20 premature branches (Fig. 3C), indicating that the ER and microtubules co-migrated to the branching axon.

### Ribosomes are in action at axon branch sites but not along the stable axon shaft

It is critical that neurons control their local environment due to their polarized and compartmentalized morphologies. To react rapidly to local requirements, specific areas of neurons may locally translate proteins(Holt et al., 2019); however, direct observation of localized protein synthesis is challenging. Our tomographic observations of axons provide direct evidence of ribosomes (Fig. 1, 2, 4, S4), particularly at axon branching sites where we found clusters of ribosomes in 29 of 63 axon branches. Ribosome clusters were found in 65% of premature axons and 37% of mature axons, indicating that ribosome clusters localized to developing axon branches. Ribosomes were also found in the filopodia at the growth cone (Fig. S6A, S6B). In contrast, we did not observe ribosomes along the tightly packed axon shafts. The visualization of ribosomes provides direct evidence that proteins are synthesized locally in the distal area of the axon, the site of dynamic cellular activities. The clustered, closely-packed ribosomes were close each other with a distance of 29.5 nm ± 3.4 nm (Fig. 4F, 4G), similar to the distance between adjacent ribosomes in a polysome (25–35 nm)(Brandt et al., 2010; Brandt et al., 2009; Mahamid et al., 2016). Therefore, clustered ribosomes likely represent a polysome in which ribosomes are arranged along a strand of mRNA, sequentially synthesizing protein. Short distances between ribosomes were observed consistently in axon branches with 70% of ribosomes falling within this short distance range (Fig. 4H, S4). This suggests that most of the ribosomes (in polysomes) in the branching axons were engaged actively in the polysome based translation, synthesizing the same types of proteins. However, 13 % of ribosomes were more than 50 nm from the nearest ribosome, too far to be considered a polysome. Although our observation does not offer to exclude if these ribosomes are in the resting state, not engaged in protein synthesis, it agrees with the notion that isolated ribosomes, termed monosomes, may be active in neurons, especially in synthesizing proteins that are only needed locally(Heyer and Moore, 2016; Holt et al., 2019). In contrast, the majority of ribosomes found at synapses are reported to form monosomes (Biever et al., 2020).

**Figure 4.**
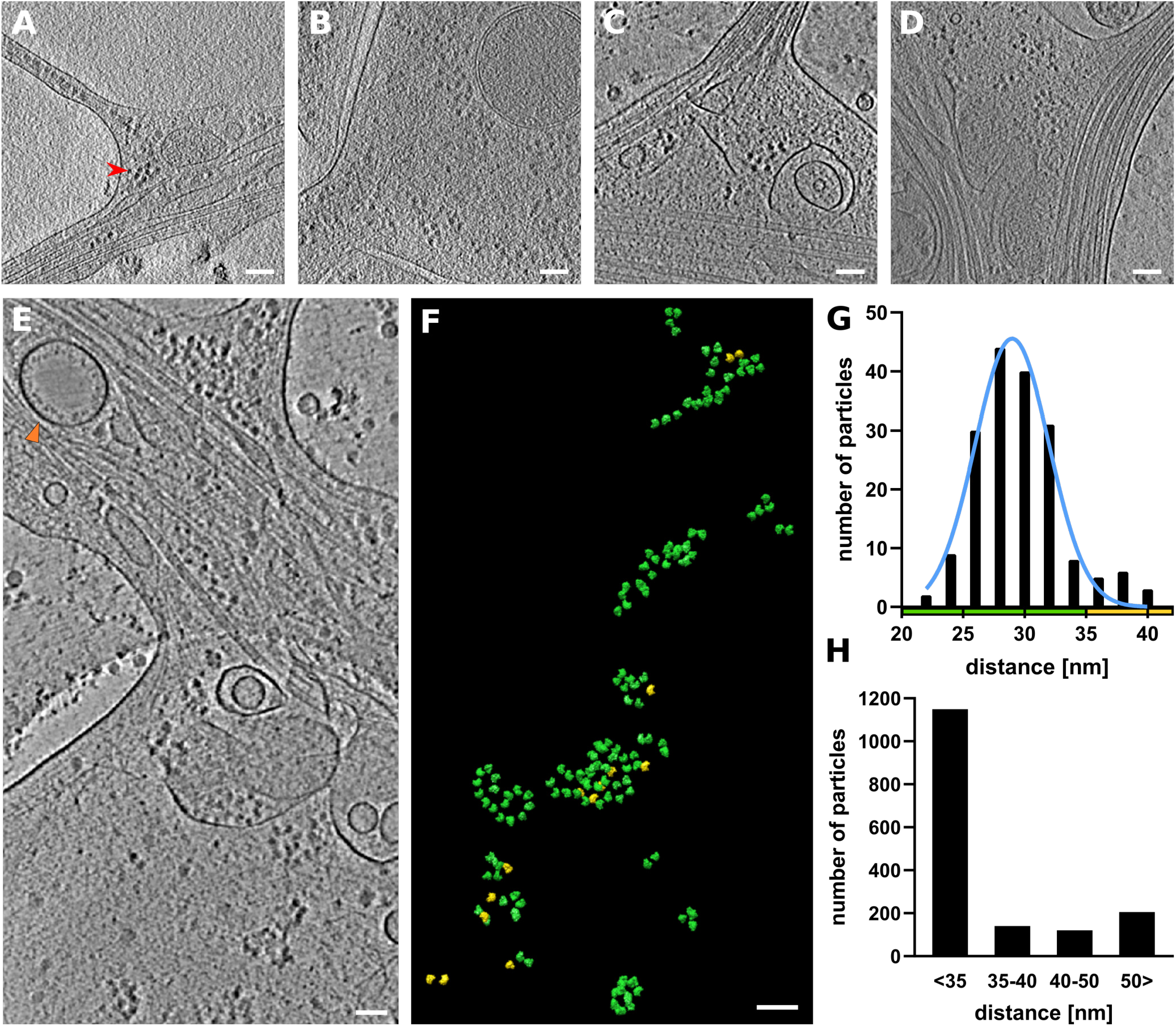
Ribosome clusters at axon branch. A)-D) Examples of clustered ribosomes in axon branches. A) and B) Ribosomes filling premature branch. Red arrowhead depicts an example of ribosome density. C) and D) Ribosomes present at mature branch site. E)-G) Analysis of distance between ribosomes. E) Slice of analyzed tomogram. (Of interest: Orange triangle shows vesicle with inner membrane densities) F) Distribution of ribosomes in the 3D volume of tomogram in E, color code by distance between ribosome particle coordinates: green < 35 nm, yellow > 35 nm. G) Distance distribution of ribosome particles in tomogram E, graph shows particles with closest neighbor distance value bellow 40 nm, N=178. H) Cumulative distance distribution for all analyzed ribosomes. N=1614. Scale bar: A-F = 100 nm.

To understand the molecular topology of ribosomes in axon branches, we computationally extracted 1614 ribosomes and performed subtomogram averaging to 38.4 nm resolution (Fig. 5A, 5B, S7). The parameters of the orientation of the individual ribosomes derived from the analysis were plotted back to the original tomograms to visualize the polysome arrangement (Fig 5D, S5). The ribosomes formed a spiral, shape resembling to the reported polysome arrangements (Myasnikov et al., 2014) (Brandt et al., 2010; Mahamid et al., 2016), indicating that they were in a polysome aligned along a strand of RNA. Although most of the observed ER in axons had no bound ribosomes regardless of their morphology, we found a few instances in which ribosomes were attached to the surface of the planar area of the ER mesh network (Fig. 5E–G). The ER-bound ribosomes adopted a polysome-like organization forming a spiral similar to those in the cytoplasm. Those ribosomes synthesize transmembrane proteins, as suggested by a cell biological analysis, but they have not been observed by ultrastructural analyses(Merianda et al., 2009). When the ER formed a thin tubular shape, there were no ribosomes attached to the membrane surface, presumably due to the high curvature. This also explains the absence of ribosomes along the axon shaft.

**Figure 5.**
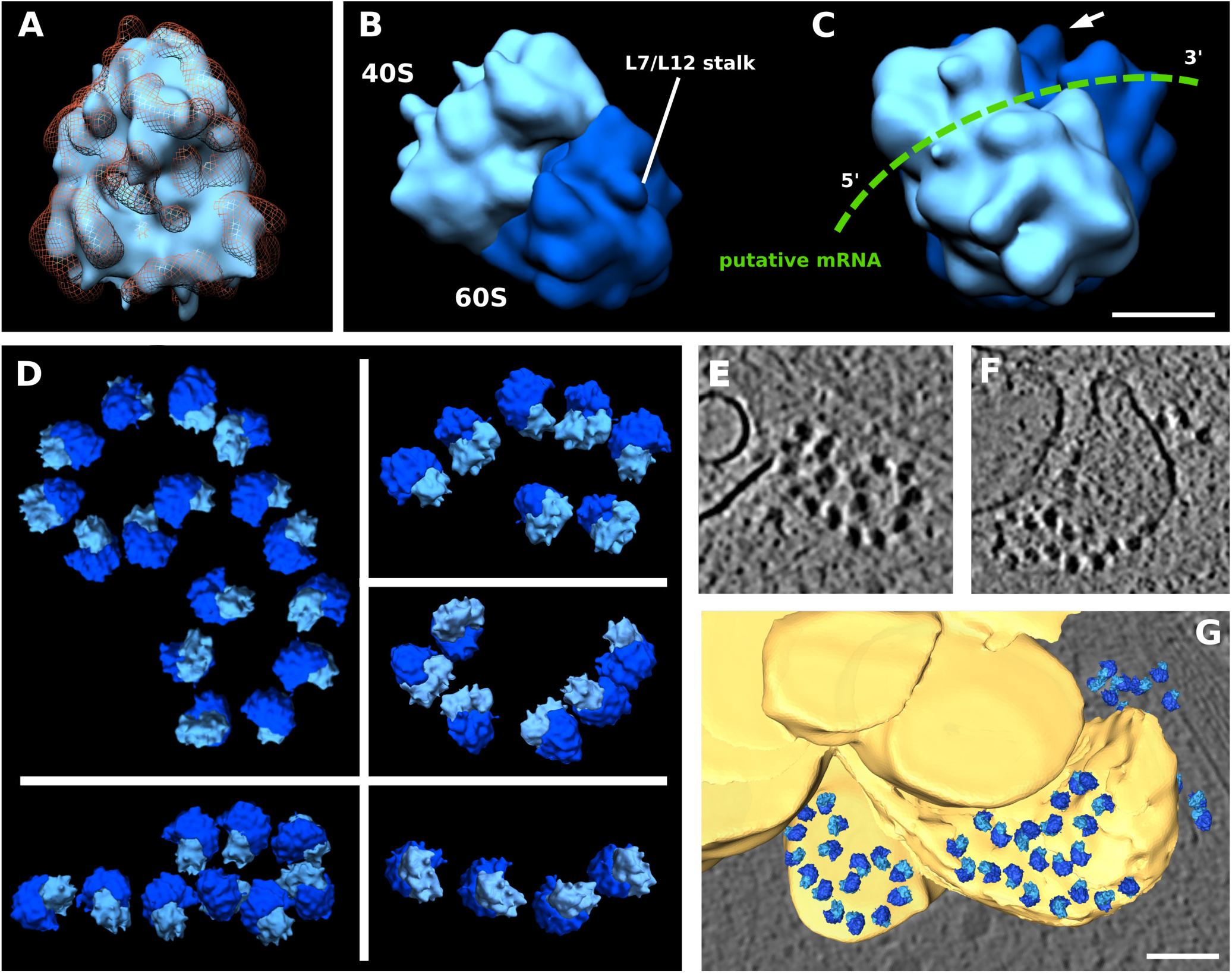
Ribosome reconstruction and polysome orientation. A) Reconstructed ribosome (light blue) fitted into the 80S ribosome volume from database (orange mesh, EMD-3420). B) Reconstructed ribosome volume with depicted 40S (light blue) and 60S (dark blue) subunits and L7/L12 stalk. C) Rotated view of ribosome from B, Putative path of mRNA depicted inn green, White arrow shows L7/L12 stalk. D) Polysomes found in various tomograms. E)-G) Ribosomes on the surface of ER. E), F) slice from tomogram, G) Segmented view for E and F. Scale bar: C = 10 nm, G = 100 nm.

## Discussion

### Molecular orchestration at the axon branch

Neuronal polarization is a unique cellular developmental process that is of critical importance. Direct observation of the process is central to gain an overview of the orchestration of the involved players, however it has been long missing. Here, using cryo-electron tomography of mouse primary hippocampal neurons and thalamus explants, we provide snapshots of mature and premature axon branches with maps of the molecular players. Axons are thin and filled with microtubule bundles, therefore, the space for cellular events is limited. That could be attributed to the fact that the mature axon is stable with a tightly packed robust cytoskeleton. At the axon branching point, the microtubule bundle bifurcates at the axon branching point, providing space for additional cellular components and dynamic cellular events. Indeed, at the branching point, we found actual evidence of ongoing cellular activities, including active mitochondria undergoing fission, active ribosomes, spreading of the ER, and fragments of actin filaments.

Our study presents observation for ribosome clusters locally concentrated at the branching point, providing direct proof of local protein synthesis at the axon branching point. In contrast, ribosomes were spread widely within less specialized cells, indicating that protein synthesis is tightly regulated within various regions of the axon. How these ribosomes are regulated and accumulate at the branching site, which is distant from the cell body in the soma, is unknown. Recently it was reported that ribosomal protein components and their coding mRNA are essential for axon branching (Shigeoka et al., 2019), giving a possible scenario that the ribosome itself may as well be remodeled locally. Identifying the steps in ribosome remodeling *in situ* at the branching point would be challenging, However, it would provide visual cues that could explain site-specific regulation of the molecular machinery that controls neuronal dynamics and the processes that lead to the establishment of a new structure like the axon branch.

### Cytoskeleton remodeling

The mechanisms that drive the rearrangement of cytoskeleton elements, microtubules and actins, at the axon and the axon branch are poorly understood. Actins are found sparsely along the axon shaft outside of microtubule bundles. Actin rings form perpendicular to the axis of the axon (Vassilopoulos et al., 2019; Xu et al., 2013) and they are predominant actin structure along the axon, although technical limitations constrain our tomographic reconstruction of these rings. At the axon branching point, abundance of short, unaligned fragmented actins accumulated, accompanied by aligned actin filaments forming filopodia-based membrane protrusion. Upon maturation of the branch, actin density decreased at the branched axon, but fragmented actins still remain at the branching point. Where are these actins originated from? Interestingly, actins are indicated to be locally synthesized during the axon branch development (Spillane et al., 2013; Wong et al., 2017). Our data on the colocalization of short actins and active polysome support the notion and bring a hypothesis that ribosomes synthesize actin as part of the machinery to build up filopodia, which is the major central process for forming cell shape.

Stabilization of the microtubules within axons requires neuronal microtubule-associated proteins (MAPs) such as Tau, MAP7, and DCX (Baas et al., 2016; Chen et al., 1992; Moores et al., 2004; Tymanskyj and Ma, 2019), but at the axon branching point, dynamic remodeling of microtubules facilitates new branch formation (Yu et al., 2008). Among the factors involved in the microtubules at the axon branch, we have previously reported a novel microtubule nucleation factor SSNA1, which localizes at the axon branching site, promotes axon branching (Basnet et al., 2018). *In vitro*, we observed a surprising microtubule nucleationthat facilitates the creation of a lattice sharing microtubule branch. Axon branching correlates with the loss of microtubule-branching activity *in vitro*, leaving open questions about the remodeling of the microtubules during axon branching. While these interesting insights are to be addressed, the assessing the microtubule bundles by electron tomography was challenging because the branching axon was too thick to allow visualization of individual microtubules within bundles. Determining how branched cytoskeleton bundles are made will be an important area for future research on axonal development.

Furthermore, we observed the ER and microtubules comigrated to the axon branching point, while the ER was rarely observed in the premature branches. The ER membrane may serve as a lipid source for the growing plasma membrane of the branching axon. The comigrating ER and microtubules may cooperate to grow stably towards branching axons, as they do in the general axon(Farias et al., 2019).

### Implications of axon branching in neural network formation

Our study provides a direct close-up view of axon branches that have a remarkable local concentration of cellular machineries, which are critical for axon development and outgrowth. This picture of the branches presents a contrast to the stable structure of the axon shaft, which serves as a rail for the transport of diverse signals and materials. It is intriguing to find a synthesis hub supporting dynamic cellular activities within the confined space of axon branches. Future studies will address the mechanisms by which the cellular machineries are recruited to and regulated at the axon branches. The process of axon branches is critical for neural network formation during the development of the nervous system. Moreover, it plays a critical role in neural homeostasis throughout the life cycle of the brain, including axon pruning during the maturation of the brain and axon regeneration after brain injury. Elucidating the mechanisms governing the formation of axon branches will not only provide insights into fundamental neuronal processes but also be the basis for understanding of neuronal circuit formation and function.

## Acknowledgements

We thank Daniel del Toro and Rüdiger Klein (Max Planck Institute of Neurobiology, Germany) for giving us advices for neuronal culture, Elena Conti, Wolfgang Baumeister, particularly, Juergen Plitzko, Ben Engel, and Marion Jasnin (Max Planck Institute of Biochemistry, Germany) for their advice and help in the initial stage of the project. The data was collected at the cryo-EM core facility at Max Planck Institute of Biochemistry, Germany and MICEF at the National Institutes of Health, USA. NM acknowledges the Max Planck Society, Boehringer Ingelheim Foundation Plus 3 Program, and the European Research Council (ERC-CoG, 724209), and the Intramural Research Program of the National Heart Lung and Blood Institute, and the National Institute of Arthritis and Musculoskeletal and Skin Diseases of National Institutes of Health, USA, for funding. NM is a recipient of the EMBO Young Investigator award.

## Methods

### Neuron cultures on EM grids

Quantifoil (R1/4 Au200, MultiA Au300, R1.2/1.3 Au300) gold EM grids were plasma cleaned for 40s and sterilized by UV light for 30 min. Grids were then coated with 1 mg/ml poly-L-lysine in 0.1 M borate buffer (Sigma-Aldrich) overnight, then washed three times in PBS and coated with laminin (Sigma-Aldrich) for 4 hours (5 μg/ml for suspension culture, 20 μg/ml for explants). Grids were washed three times with PBS and covered with neurobasal/B27 medium and incubated at 37 °C.

Primary embryonic mouse neurons were prepared as either dissociated hippocampal cultures or as a tissue explants from thalamus. Neurons were prepared from embryonic day 15.5 (E15.5) mice. Dissected hippocampi were placed into cold HBSS (HBSS supplemented with 1x HEPES, 1x Glutamax, 1x Pen/Strep) media treated with trypsin incubating at 37 °C for 16 min followed by washing with HBSS with FBS and then neurobasal/B27 medium followed by trituration. Thalamus tissue was placed into neurobasal/B27 medium and explants were prepared by cutting thalamus into small pieces which were incubated at 37 °C for 30 min. 11 of the observed cryo-EM images of hippocampus neurons were from the effort of transducing microtubule binding protein SSNA1. The transduction rate was low and furthermore microtubule organization was not assessed in this study. The other components did not show any notable differences to the extent of our experimental evaluation. The dissociated cells were plated on coated EM grids at a concentration 150,000 cell/ml and incubated at 37 °C in 5% CO2. For the explants, they were placed on EM grids covered in neurobasal/B27 medium. The culture medium was NeuroBasal (Invitrogen) supplemented with 1x Glutamax, 1x B27 serum and 1x Pen/Strep (neurobasal/B27 medium). Half of culture media was changed at DIV 1 (day in vitro). Cell cultures were grown for 6-10 days. Then EM grids with neuron cultures were manually vitrified in liquid ethane using home-made plunger or vitrobot (ThermoFisher Scientific).

### Cryo-ET data collection and processing

Cryo-electron tomography data were collected on Titan Krios (ThermoFisher Scientific) with a Gatan Quantum 967 LS and K2 Summit direct detector with an option of phase-plate with an acceleration voltage of 300 kV. 63 tilt-series were collected using phase-plate option and 54 tilt-series were collected without phase-plate with defocus range between −3.5 μm to −5 μm. Tilt series were collected −60° to 60° with 2° angular increment using does symmetric scheme using Serial-EM software (Hagen et al., 2017). The total electron dose was around 90 e-/Å and the nominal magnification was 26,000 x, corresponding to the final pixel size of 5.46 Å. Images were taken in super-resolution mode as ten-frame movies and the movie frames were aligned and combined and dose filtered using in-house frame alignment software with implemented MotionCor2. A total of 117 tilt-series were assessed for this study including 52 tilt-series of thalamus explants and 65 tilt-series of hippocampus neurons.

Individual images of the collected tilt-series were assessed manually and low-quality images at the high tilt angle were removed from the dataset. Tilt-series were filtered according to the cumulative radiation dose (Grant and Grigorieff, 2015) and aligned on the basis of the patch tracking algorithm using the IMOD ETOMO package (Kremer et al., 1996). Tomograms were reconstructed from aligned stacks using weighted back-projection in IMOD. Tomograms were further 4x binned, resulting in pixel sizes of 21.8 Å. Tomograms were denoised by edge enhancing diffusion (bnad command in Bsoft(Heymann et al., 2008)).

### Tomogram segmentation

Tomograms were manually segmented using Amira software. Additionally when applicable, membranes were segmented automatically using deconvolution filtering(Tegunov and Cramer, 2019) and a tool for membrane segmentation TomoSegMemTV(Martinez-Sanchez et al., 2014). Microtubules were segmented manually in IMOD software. Measurements of mitochondria length (44 mitochondria) and ER tube diameter (19 for thinnest ER measurements and 29 for ER tube diameter) were measured manually in IMOD. The data obtained were represented using box and whiskers graph, where each dot represents measurement for individual mitochondrion or ER and the horizontal line in the graph indicates the median of the distribution. Data was plotted using Prism software.

### Subtomogram averaging of ribosomes and distance analysis

1614 ribosome particles from 11 unbinned tomograms were picked using IMOD 3dmod software. The picked coordinates were then transferred into Dynamo a subtomogram averaging software package (Castano-Diez et al., 2012) and the ribosome averages were calculated as follows. The initial template used for the alignment was low pass filtered to >100 Å (EMDB-5224). At this resolution, only general shape and size of the ribosome was visible. The initial alignment was done using standard global settings from Dynamo and the search space and angular increments were then gradually decreased for subsequent refinement. During refinement, the subtomograms were split into odd and even half-sets and Dynamo’s adaptive bandpass filtering was performed in order to avoid overfitting and estimate the attained resolution. The attained resolution was estimated by comparing the FSC of separately computed averages from odd and even half-sets. A bandpass filter was then applied in next iteration based on this estimation. Distances between ribosome particles were calculated from refined coordinates from subtomogram averaging runs using Python3 (numpy and scipy libraries). For each particle, the closest neighboring distance was plotted into the distance distribution histogram and fitted with non-linear Gaussian curve in Prism software. The orientation of ribosomes was assessed by placing reconstructed ribosome volume into tomograms in Chimera using coordinates and alignment parameters derived from subtomogram averaging.

---

**Figure S1.**
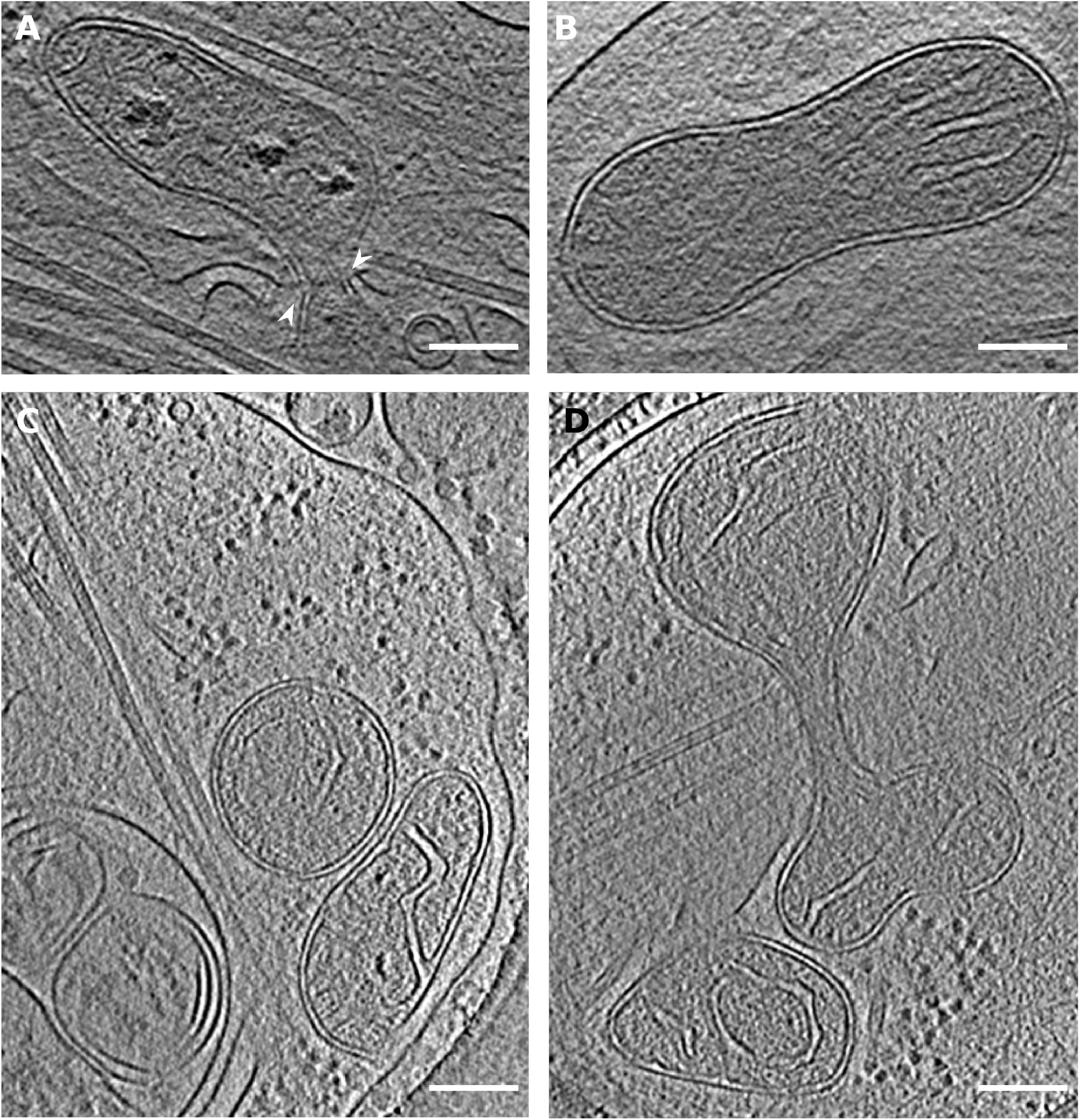
Examples of mitochondria found at axon branch. A) Mitochondrion undergoing fission facilitated by ER. White arrowheads depict ER pinching at mitochondria membrane. B) Possibly dividing mitochondrion. C) Examples of different sizes of mitochondria. D) Mitochondrion presumably undergoing fission. Scale bar: A-D = 100 nm.

**Figure S2.**
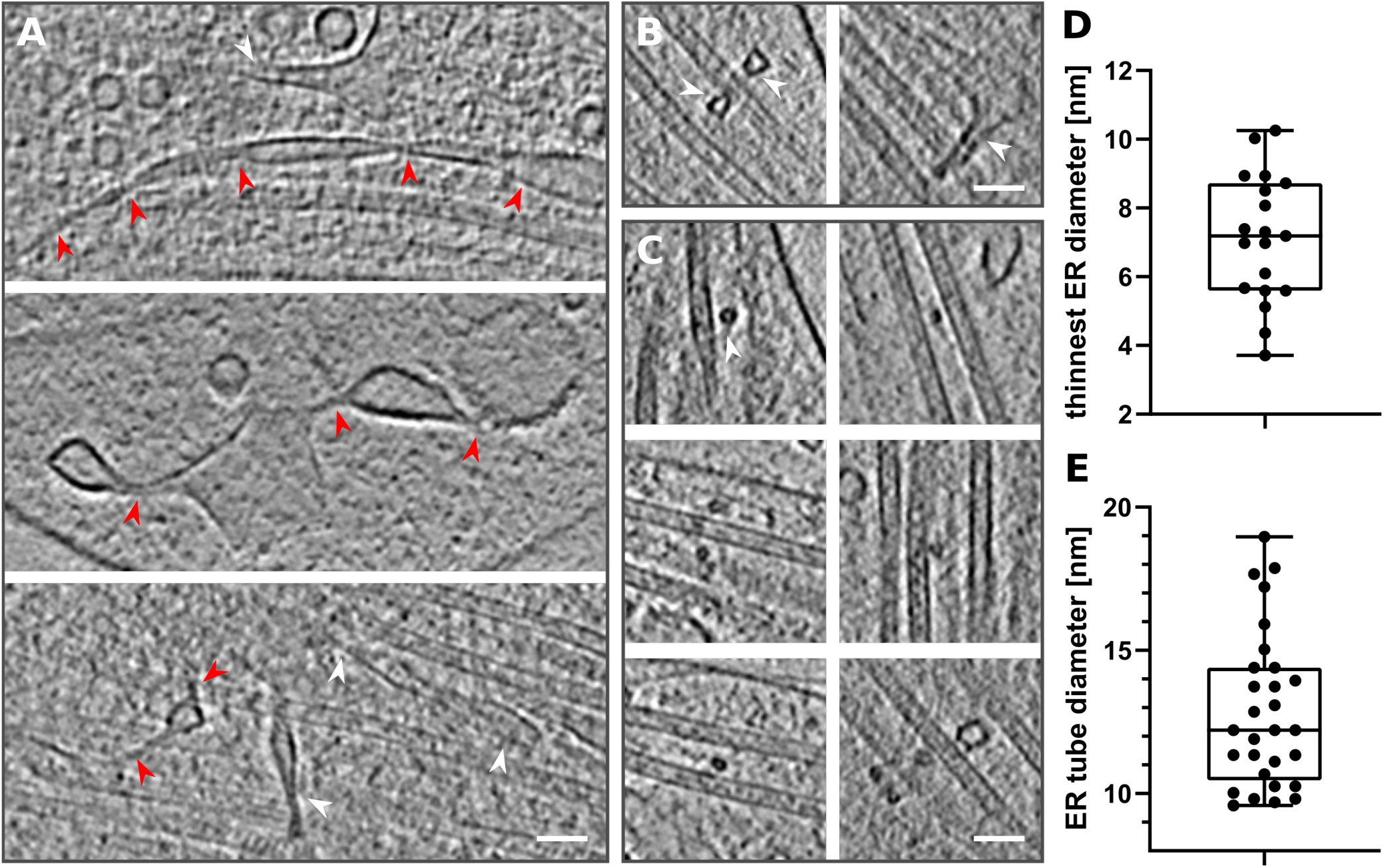
Thin ER tubes in axon branches. A) Examples of ER tube reaching extremely thin diameters. Red arrowheads follow continuous ER tubes, White arrowheads depict different ER. B) ER (white arrowheads) wrapping around microtubules. C) Cross-section of ER tube (white arrowhead) wrapping around microtubules. D) Diameter of thinnest ER tubes found in tomograms, median diameter: 7.19 nm, N=19. E) Diameter of ER tube wrapping around microtubule, median diameter: 12.21 nm, N= 29. Scale bar: A-C = 50 nm.

**Figure S3.**
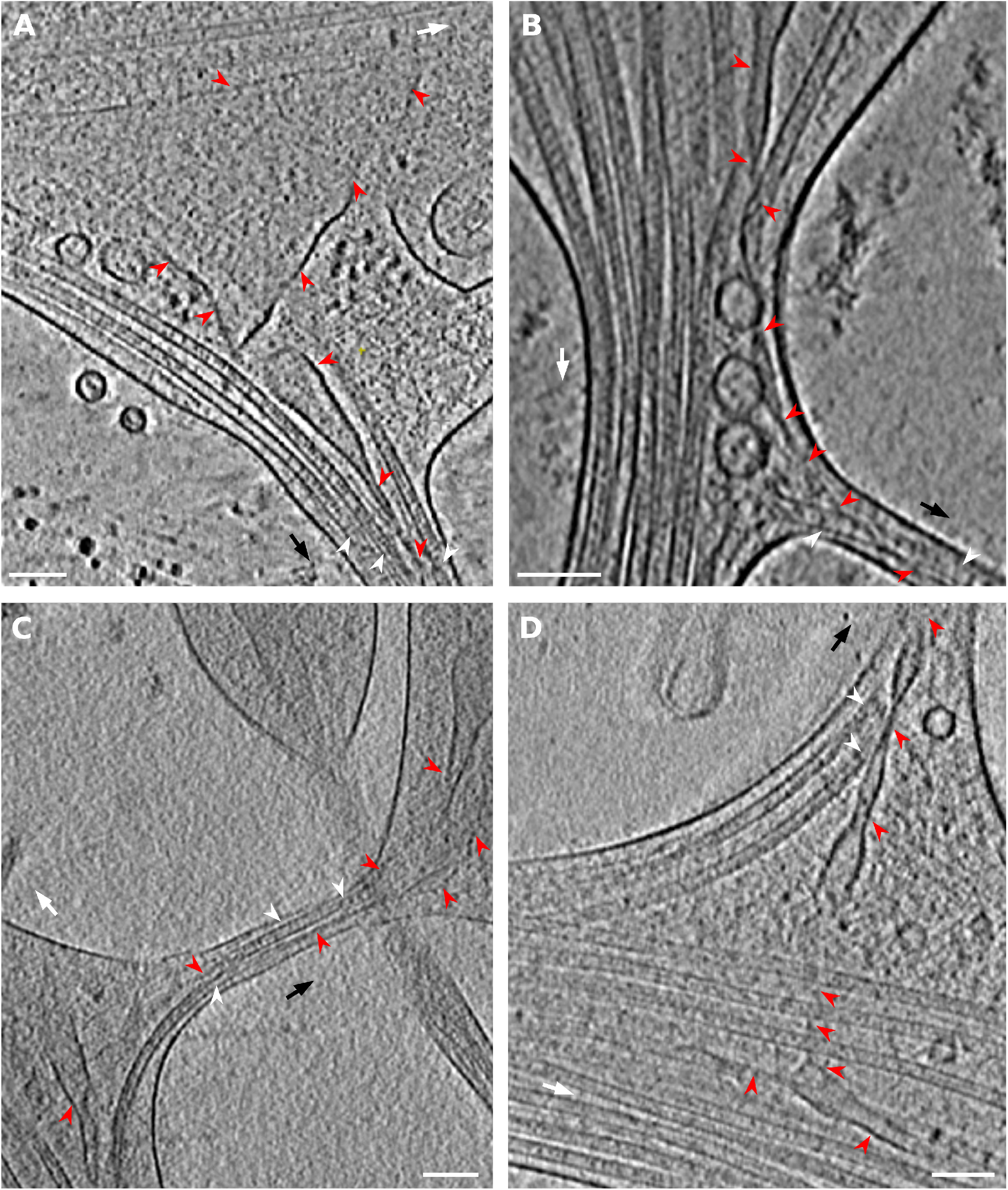
ER enters axon branches together with microtubules. A)-D) Examples of ER tube present in axon branch together with microtubules, red arrowheads – ER, white arrowheads – microtubules, black arrow shows direction of new branch, white arrow points in the direction of main axon growth. Scale bar: A-D = 100 nm.

**Figure S4.**
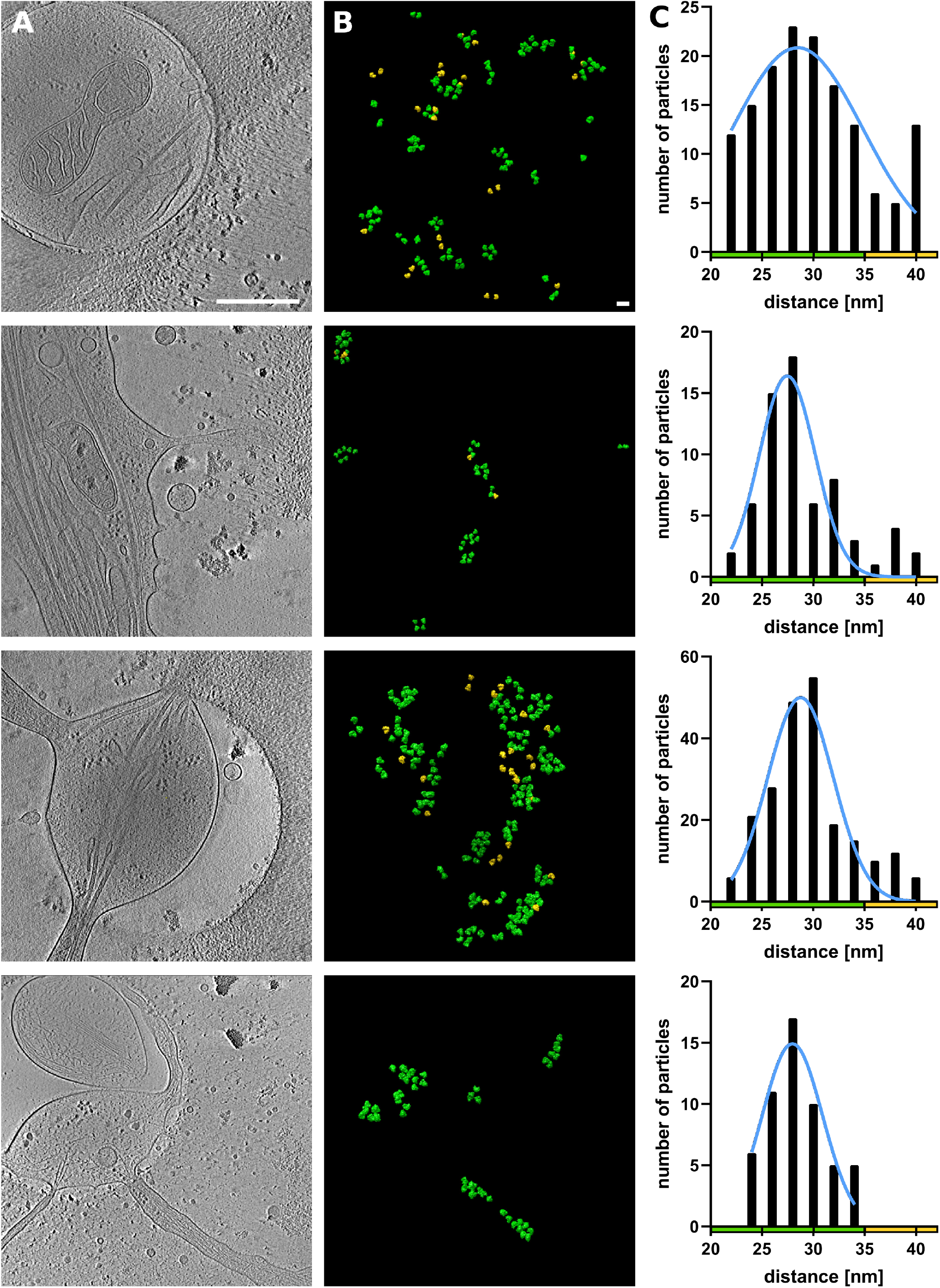
Visual representation of the ribosome distance distribution. A) Slice from tomogram. B) Ribosome particles placed into the tomogram volume in color representation according their closest neighbor distance, green < 35 nm, yellow > 35 nm. C) Graph representing the distance distribution between clustered ribosomes. Scale bar: A = 500 nm, B = 50 nm.

**Figure S5.**
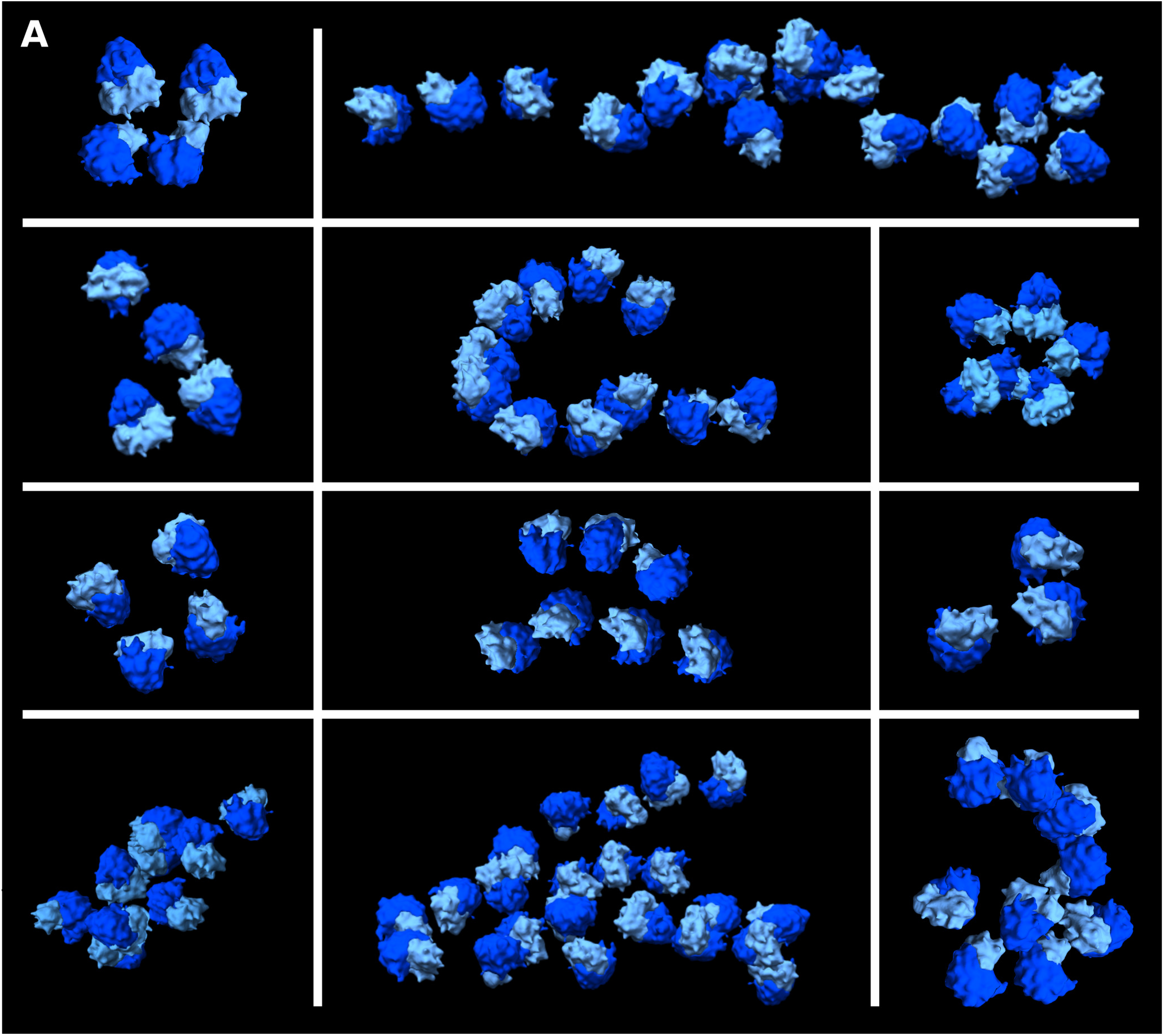
Clustered ribosomes form polysomes. Examples of strings of ribosomes found in axon branches forming polysomes.

**Figure S6.**
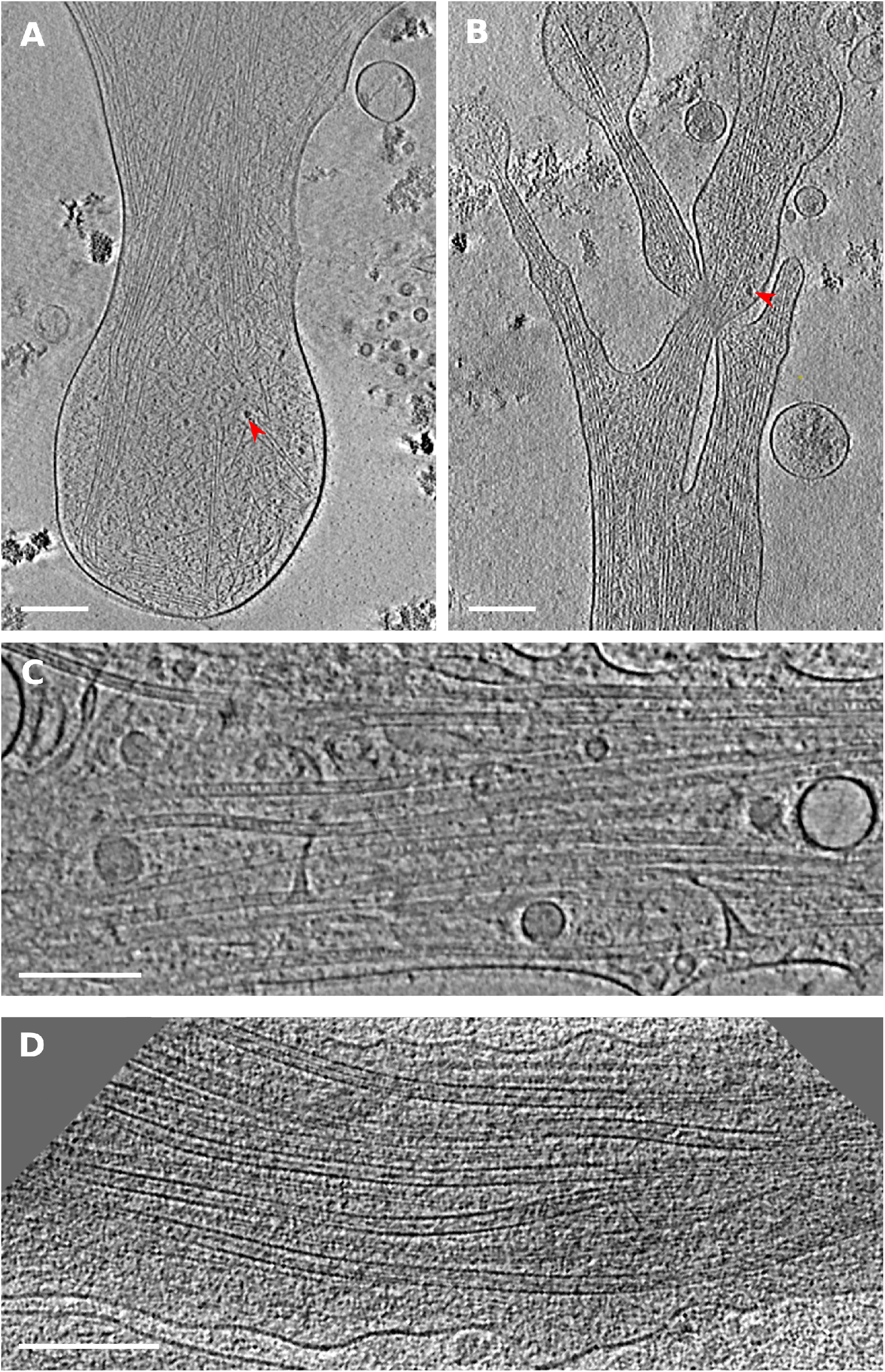
Growth cone and axon shaft. A), B) Slices of tomograms showing growth cone filled with actin, red arrowhead depicts ribosome. C) Axon shaft packed with bundled microtubules, thin ER tubes and vesicles. Scale bar: A-C = 200 nm.

**Figure S7.**
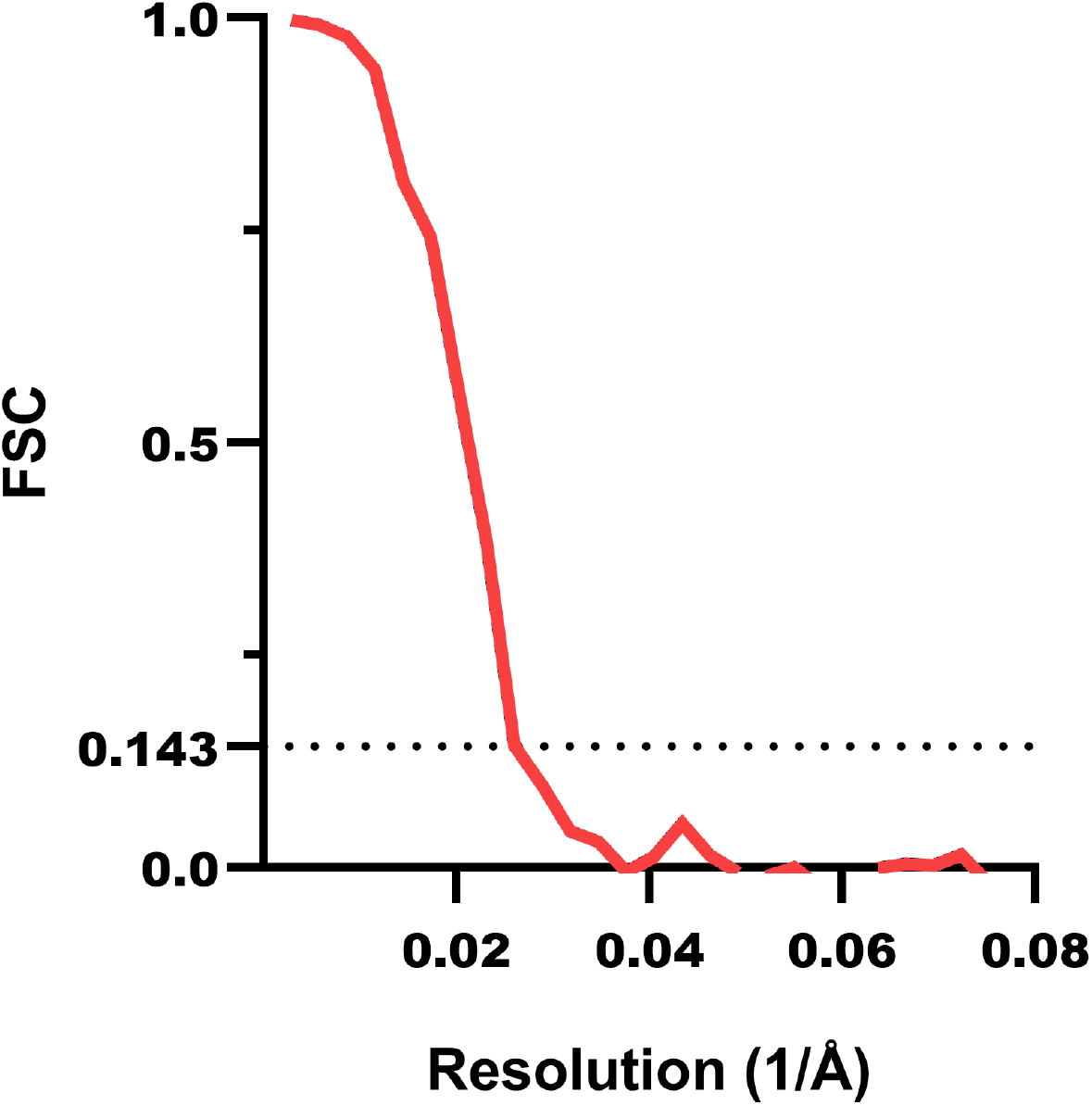
Ribosome reconstruction resolution estimation, FCS curve.

